# Cyclic Microchip Assay for Measurement of Hundreds of Functional Proteins in Single Neurons

**DOI:** 10.1101/2021.06.06.447288

**Authors:** Liwei Yang, Avery Ball, Jesse Liu, Tanya Jain, Yue-Ming Li, Jun Wang

**Affiliations:** Multiplex Biotechnology Laboratory, Department of Biomedical Engineering, Stony Brook University, Stony Brook, NY 11794; Chemical Biology Program, Memorial Sloan Kettering Cancer Center, New York, NY, USA; Programs of Neurosciences, Weill Graduate School of Medical Sciences of Cornell University, New York, NY, USA; Programs of Pharmacology, Weill Graduate School of Medical Sciences of Cornell University, New York, NY, USA

## Abstract

Proteins are responsible for nearly all cell functions throughout cellular life. To date, the molecular functions of hundreds of proteins have been studied as they are critical to cellular processes. Those proteins are varied dramatically at different statuses and differential stages of the cells even in the same tissue. The existing single-cell tools can only analyze dozens of proteins and thus have not been able to fully characterize a cell yet. Herein, we present a single-cell cyclic multiplex *in situ* tagging (CycMIST) technology that affords the comprehensive functional proteome profiling of single cells. It permits multiple, separate rounds of multiplex assays of the same single cells on a microchip where each round detects 40-50 proteins. A decoding process is followed to assign protein identities and quantify protein detection signals. We demonstrate the technology on a neuron cell line by detecting 182 proteins that includes surface makers, neuron function proteins, neurodegeneration markers, signaling pathway proteins and transcription factors. Further study on 5XFAD mouse, an Alzheimer’s Disease (AD) model, cells validate the utility of our technology which reveals the deep heterogeneity of brain cells. Through comparison with control mouse cells, the differentially expressed proteins in the AD mouse model have been detected. The single-cell CycMIST technology can potentially analyze the entire functional proteome spectrum, and thus it may offer new insights into cell machinery and advance many fields including systems biology, drug discovery, molecular diagnostics, and clinical studies.

## Introduction

Recent advancement of single-cell omics technologies has revolutionized the study of complex biological systems and has significantly impacted on precision medicine^1, 2^. A scale at the omics level is necessary since a typical animal cell contains numerous proteins, transcripts and other molecules that participate in normal functions^3, 4^. Propelled by the broadly available sequencing tools, single-cell transcriptomics has been frequently applied in many human diseases to reveal the genome-wide gene expression among individual cells^5, 6^. Proteins, however, are not amplifiable like DNAs, and thus protein analysis in single cells has not reached the scale comparable to transcriptomes. Proteins largely represent cell functions and biomarkers for disease diagnosis and cell type classification^7^. Functional proteins (hundreds as known) predominate for representing cell identity, drug target, clinical biomarkers, signaling networks, transcriptional factors, functional readouts of proliferation, cell cycle status, metabolism regulation and apoptosis makers^7, 8^. Particularly in disease pathology and signal pathway analyses, single-cell functional proteomics is irreplaceable, while the correlation of gene expression and protein expression is generally ~0.4 and almost no correlation for low expression proteins (particularly for signaling proteins)^9, 10, 11^. Furthermore, protein-level diagnosis desirably offers more direct implication of pathogenesis and potential drug targets^8^.

A variety of multiplex strategies have been introduced to achieve the measurement of dozens of proteins per cell, but they are still far from functional proteome level yet ^4, 12^. While conventional fluorescence flow cytometry can even detect 17 colors simultaneously^13^, it is barely used in practice due to high cost of instrumentation and reagents. Mass cytometry, a technique utilizing transition element isotopes as labelling tags instead of fluorophores, offers a higher multiplexity of ~40s in single-cell analysis^14^. Similarly, multiplexed ion beam imaging by time of flight (MIBI-TOF) leverages isotope tagging in immunohistochemistry to achieve high-resolution imaging of 36 proteins for tumor specimens^15^. Another strategy is to reiteratively stain cells with fluorophore tagged antibodies for a few cycles to overcome the limitation of fluorescence spectrum overlapping ^16, 17, 18, 19, 20, 21^. The same cells are imaged in each cycle and are registered, so a multiplexity of 60s-80s is achievable^18^. This is a convenient, albeit laborious, technique that does not require special instruments. Nevertheless, for all the immunohistochemistry-based techniques, the high multiplexity is associated with the high risk of ‘parking’ issue due to limited room for binding of many antibodies to protein clusters^22, 23^. A cell is packed with macromolecules, cytoskeleton and organelles, and therefore it cannot accommodate too many antibodies that are normally large molecules. These factors limit the practical use of multiplexed assays at less than 50 proteins per batch^24^. The other emerging methods based on mass spectrometry and microchip western blotting can potentially overcome the bottleneck of multiplexity, but they are still not practically useful yet for routine high-multiplex assays in single cells^25, 26^.

DNA barcoding on microarrays is a robust genome-wide assay that can analyze thousands of molecules simultaneously^27^, although it is not at the single-cell level yet. To transform its utility in protein assays, barcoded DNAs are tagged with antibodies where the antibodies recognize targets and DNAs confer signal^23, 28, 29^. A plethora of strategies can be adopted for signal amplification, enhancement and multiplexed detection with the aid of barcoded DNAs. The signal readout can rely on PCR^30^, fluorescence imaging^31, 32^, NanoString^33^, and even next-generation sequencing^29^, while their multiplexity is still limited by antibody labeling. Miniaturized DNA barcode array enables the quantification of up to 35-40 proteins in single cells when the array is combined with a microfluidic chip for single-cell manipulation^34^. Further improvement of multiplexity is limited by the size of DNA arrays which are normally 20 μm for each^35^. In our previous work, we have developed a multiplex *in situ* tagging (MIST) array technology on DNA encoded microbeads for single-cell cytokine assays^36, 37^. The small size of microbeads holds great potential for functional proteomic assays in single cells since MIST has increased the assay capacity by 100 times.

Herein, we report a single-cell cyclic multiplexed *in situ* tagging (CycMIST) platform that can analyze hundreds of protein targets in single cells with high throughput and high sensitivity using a common imaging platform. The high protein content is achieved by the multi-round labeling and multi-cycle decoding process based on the CycMIST platform. Each single cell in a microchip microwell can be stained for 4 rounds by a cocktail of UV-cleavable DNA barcoded antibodies, and for each staining the DNA oligos can be released by UV irradiation and captured by a MIST array. The decoding process on a MIST array permits identification of up to 50 types of released DNAs at one time. Thus, the whole process overcomes the typical parking issue related with high multiplexity while the total multiplexity can increase by 40-50 for each round. We have thoroughly validated the technology and characterized its performance. As a proof-of-concept study, the CycMIST has been applied to differentiated mouse neuroblastoma Neuro-2a (N2a) cells for analysis of 182 proteins that include surface makers, neuron function proteins, neurodegeneration markers, signaling pathway proteins and transcription factors. An additional study on tissue from the mouse pre-frontal cortex found that CycMIST can profile the functional proteome of individual brain cells and distinguish molecular features between WT tissue and Alzheimer’s Disease (AD) tissue from a 5xFAD mouse model. The CycMIST technology should be broadly applicable to various complex diseases in the future for mechanistic studies at the single-cell functional proteomics level.

## Results

### Workflow of single-cell CycMIST technology

The CycMIST platform involves multi-round staining/dissociation and multi-cycle decoding processes for multiplexed single-cells protein profiling (Fig. 1a). Each round can detect up to 50 proteins simultaneously for a single cell, and the decoding process for each round is to assign the protein ID to the detected protein signals. Specifically, cells are loaded and fixed in PDMS microwells first before immunostaining by a mixture of complementary DNA (cDNA)-antibody conjugates. Each conjugate contains two UV cleavable linkers between the cDNA and the antibody, and one end of the cDNA has a biotin moiety. A MIST array is mated with the PDMS microwells with assistance of a clamp to completely seal the microwells. UV light exposure through the device releases the cDNAs, which are specifically captured by the ssDNAs on the MIST array by hybridization. Then the MIST array is separated from the PDMS. Those bound antibodies on the cells are dissociated by a regeneration buffer, initiating the next staining round by another conjugates panel for different targeting. Each round of staining/dissociation generates a MIST array with fluorescence signals corresponding to protein abundance which is denoted as protein signal. Decoding on the MIST array requires multiple cycles which are dependent on the number of targets (multiplexity). It relies on the unique, consecutive color change on the same microbead to recognize the DNA sequence, or the detected protein, by this microbead, since each DNA sequence corresponds to only one type of protein target as is predesigned (Table S1), and all DNAs are orthogonal to each to each other. To track the same microbead from protein detection to decoding experiments for all ~25 million microbeads of a MIST array, the images from protein signal and the images of decoding are well aligned by a Matlab program. Notably, the MIST array could be decoded beforehand or after protein detection. In the current report, we demonstrate the detection of 182 proteins through 4 rounds of staining/dissociation and 4 cycles of decoding process with 3 colors dye. Higher multiplexity can be simply achieved by more rounds of staining/dissociation with up to 50 proteins for each, and thus hundreds of proteins can be detected for single cells. Theoretically, an unprecedented level of multiplexity (*K* × *M^N^*) can be achieved through *K* rounds of cellular staining, in combination with *N* decoding cycles by *M* color types.

**Figure 1.**
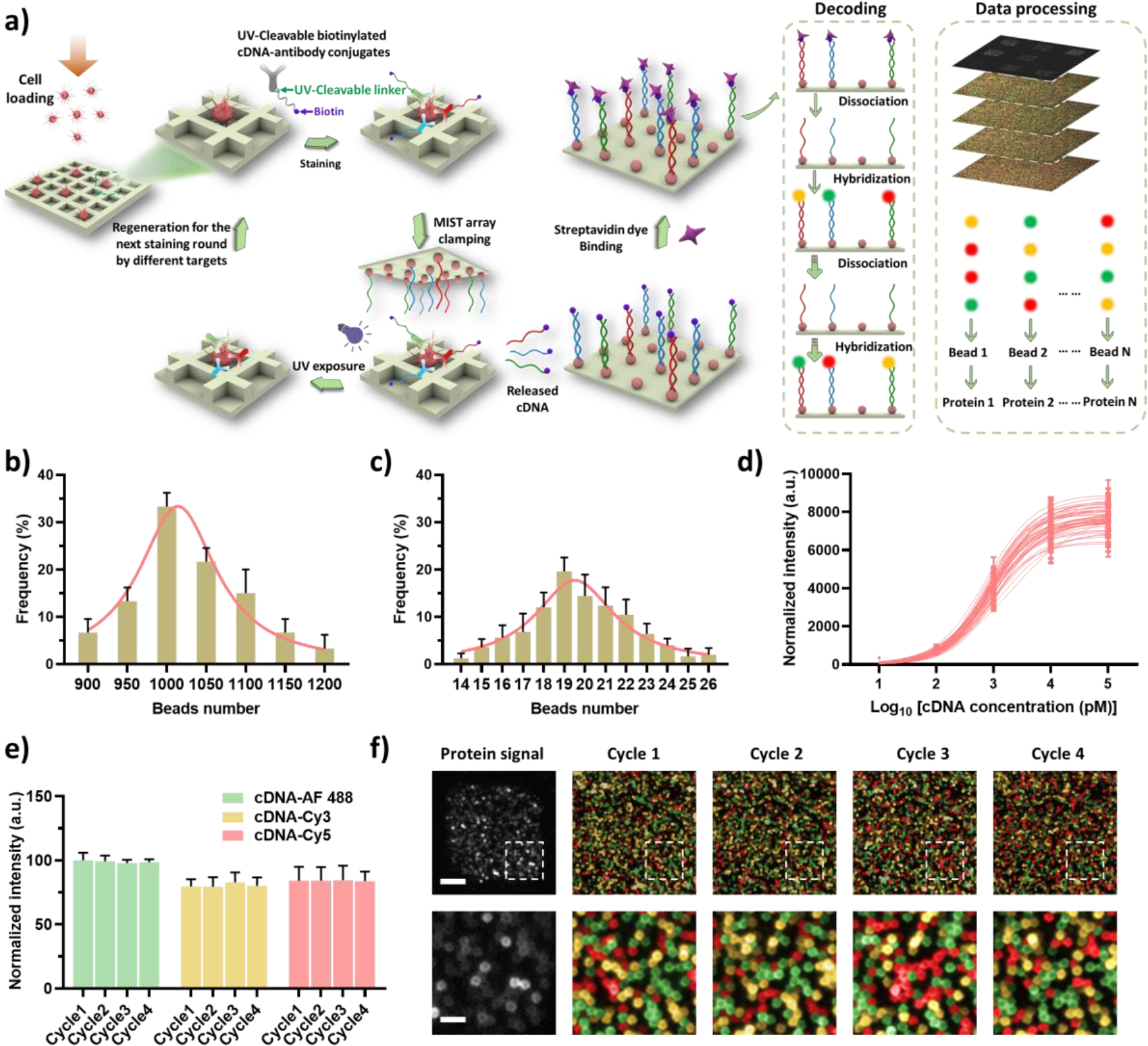
Overview of CycMIST technology for single-cell functional proteome analysis. a) Schematic illustration of the CycMIST process to analyze multiple proteins through the MIST microbeads array. b) Distribution of the number of ssDNA-coated microbeads on each 75 μm × 75 μm area of a MIST array that is corresponding to a PDMS microwell. c) Distribution of the number of same kind ssDNA-coated microbeads on a microwell area. d) Characterization of the CycMIST sensitivity by varying the concentrations of 50 biotin-cDNAs on the MIST array. This is the same procedure in single-cell protein detection experiments except cell loading and conjugate binding. e) Consistency of fluorescence intensities for 4 decoding cycles and for 3 fluorescent color dyes. f) Sample images of multiplexed assay of 50 proteins from a single cell by CycMIST and the 4 decoding cycles images. The greyscale images are protein detection result, and the color images are the decoding cycles from cycle 1 to cycle 4. The bottom panel is the zoom-in images from the squares in the up panel. Scale bar: 20 μm (up panel); 5 μm (bottom panel).

### Characterization of high-density and high-sensitivity MIST array in detection

The MIST array is comprised of a monolayer of 2 μm microbeads in an area of 1 cm by 1 cm, which was facilely fabricated by attaching the DNA-coated microbeads to a pressure-sensitive adhesive tape covered on a plain glass side^37^.This is a super-compact microarray that contains ~25 million DNA coated microbeads of 50 different types (Figure S1). Each section of 75 μm × 75 μm imprinted by the PDMS microwells encloses 1012±74 ssDNA-coated microbeads in average (Figure 1b). The microbeads with the same type of ssDNA have 19 ± 4 copies per array (Figure 1c), and this copy number variation of microbeads has been demonstrated to have negligible effect on the reproducibility and quantification of protein detection in our previously reported work^36^.

The orthogonality of the 50 DNA pairs was fully examined, which shows less than 1% cross reactivities between any of noncomplementary pairs (Figure S2). The sensitivity of those DNAs on the MIST array has been also quantified through hybridization of cDNA-dyes (Figure 1d). The sensitivity has been significantly improved by the current coating method where poly-L-lysine mediates the DNA absorption on the microbeads^37^. It is found that all DNAs perform similarly in capturing their cDNAs on the array. The averaged limit of detection (LOD) for all the 50 types of DNA-microbeads is found to be around 4 pM. In consideration of the small-size microwells at 225 pL, theoretically ~542 cDNA copies released from antibodies binding to a single cell can be detectable.

The high-quality of the MIST array ensures high fidelity of 4-cycle decoding process to assign the right colors to each microbead. In each cycle, a cocktail of cDNAs-dye is applied to the MIST array where 3- color fluorescent dyes are used in this report. The signal-to-noise ratio of cDNA-dye labeled microbeads is generally above 5.5, and <2.3% microbeads have been incorrectly assigned with colors in the decoding test. Multiple cycles of hybridization and dissociation has no noticeable influence on the freely accessible DNAs on the microarray (Figure 1e), which implies even more cycles possibly available when higher multiplexity is demanded. Figure 1f shows typical resultant images of the protein detection and decoding on microbeads for the detection region. Protein signals are confined within a 75 μm × 75 μm area, and the varied brightness indicates different protein expression levels. In the same region, successive decoding results of 4 cycles are shown in 3 colors for Alexa Fluor 488 (green), Cy3 (yellow) and Cy5 (red) channels, where each microbead exhibits an ordered color change over the cycles. The microbeads with the same identities are grouped, and their fluorescence intensities are quantified and averaged to generate the protein expression profile for a cell.

### Robustness of UV cleavage and reiterative staining of cDNA-antibody conjugates

The quality of the cDNA-antibody conjugates is critical to the CycMIST assays. Testing on a few crosslinking methods results in the selection of Azido/DBCO click chemistry due to high conjugation yield and convenience (Figure S3). To increase cleavage efficiency, two UV-cleavable nitrobenzyl groups are used instead of one in the similar conjugations previously published^30, 33^. Each synthesized conjugate was purified by fast protein liquid chromatography (FPLC) and validated by a Nanodrop spectrum. Thorough study found the selection of purification process significantly affects immunofluorescence staining result. Centrifugation by 100 KDa filter, which is a typical method adopted by the field, still left a non-trivial number of free DNAs in the solution as our FPLC result showed. Even the conjugates within the peak in Figure S3 performed quite differently. Thus, we only select the top 2/3 of the conjugate peak for all following experiments. The degree of labeling (DOL) was measured to be 2.2 to 4.1 cDNAs per antibody across different conjugates. This DOL is normally deemed as the optimal one for antibody labeling.

Eight conjugates for targeting tubulin, Lamin B1, CREB, VGLUT2, MAP2, phospho-GSK3, CD24 and CD86 that covers housekeeping protein, nucleus proteins, neuronal differentiation markers, phosphorylated protein, and cell membrane proteins have been validated by immunofluorescence staining. Some in the panel have known high expression levels, and some should have no expression. In this test, the end of cDNA was tagged with a Cy5 dye instead of biotin for each conjugate. The staining result is compared with that when secondary antibodies tagged with Alexa Fluor 488 dye were used. Figure 2a and Figure S4 show that the expected compartments of the cells are stained for all the conjugates, and the staining results by both methods are consistent. Next, we optimized UV cleavage conditions and assessed the UV-cleavage performance. With our portable UV light, cells lose fluorescence completely within 15 min exposure (Figure 2b and 2c). This relatively long-time exposure should be related with UV light penetration through the thick plastic plate used for clamping and also through a layer of PDMS. Nevertheless, the cleavage has been highly efficient that no visible fluorescence is observed after UV irradiation (Figure 2c and Figure S5). The quantified intensities for single cells in Figure 2d further exhibit >97.4% of fluorescence decrease upon UV cleavage while the pre-cleavage intensities meet the expectation for those proteins.

**Figure 2.**
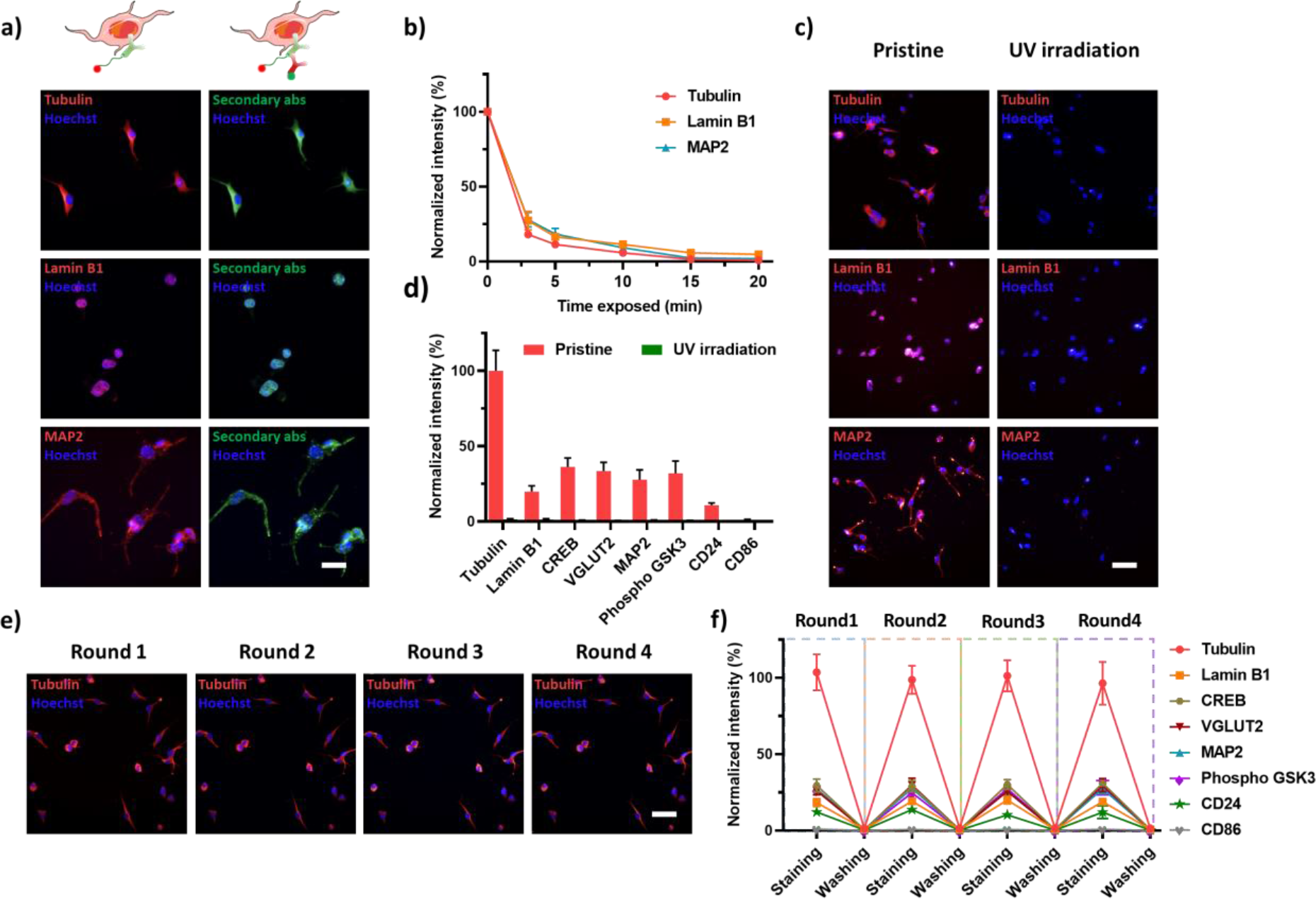
Performance of multi-round conjugates labeling and cDNA release. a) Colocalization of cDNA-antibody conjugates immunofluorescence staining labeled with fluorophore Cy5 on cDNA (left) and labeled with Alexa-488 tagged secondary antibody (right). Scale bars: 20 μm. b) Efficiency of UV cleavage over time for three cDNA-antibody conjugates. c&d) Measurement of binding location and UV cleavage efficiency for eight conjugates. Normalized intensity in d) show ~97.4% cDNA being released after 15 min UV exposure. All fluorescence intensities of conjugates staining were normalized by tubulin intensity. Scale bars: 50 μm. e&f) Reproducibility of 4 consecutive rounds of conjugates staining, releasing, and re-staining with the same cDNA-antibody conjugates. All fluorescence intensities of conjugate staining were normalized by tubulin intensity. Scale bars: 50 μm. Error bars are within symbol size if not shown.

The reiterative staining procedure has been optimized so that the same cells can be stained for multiple rounds. With this process, the antibody binding of a later round will not be affected by the previous round, which resolved the parking issue when a large panel of antibodies is used. After each staining and imaging, the cDNA-antibody conjugates were dissociated by a regeneration buffer before re-staining by another cocktail of conjugates. Our result shows that the conjugates are completely removed from the cells, which is confirmed by labeling of secondary antibodies (Figure 2e, 2f and Figure S6). Re-staining of the same cells exhibits the similar fluorescence intensities for each of the 8 conjugates with <6.5% variation. This high fidelity suggests no reduced immunogenicity of cells in the multi-round procedure, which lays the foundation for the high multiplexity of CycMIST.

### Multiplexed single-cell CycMIST for measurement of 182 proteins

The performance of the CycMIST for single-cell analysis has been investigated. Cells were attached and fixed to a Pluronic^®^ F127-coated PDMS microwells chip. Approximately only 0-2% cells were lost in each round of the experiment (Figure 3a), which were recorded by imaging. The area of lost cells on the MIST array was excluded in data processing. The experimental protocol has been optimized so that the signals detected for each microwell area on the MIST array are pertinent to a unique cell (Figure S7). The 8-panel proteins are used to validate the single-cell assays. Figure 3b displays the linear increase of fluorescence signals with incremental cell numbers (0, 1, 2, 3 and 4 cells). This proves true signals being detected from single cells. To validate the reproducibility of the technology, the same 8-panel proteins were analyzed for 4 rounds on the same cells in microwells. The detected signals are highly consistent with the bulk assay where tubulin is still the highest and CD86 is the lowest (Figure 3c). The variation of detecting the same protein across different rounds is only 8.6%±1.5% by the single-cell CycMIST technology. Thus, every round of the detection can be deemed as a separate multiplexed assay, and the multiplexity can be efficiently increased by more round numbers.

**Figure 3.**
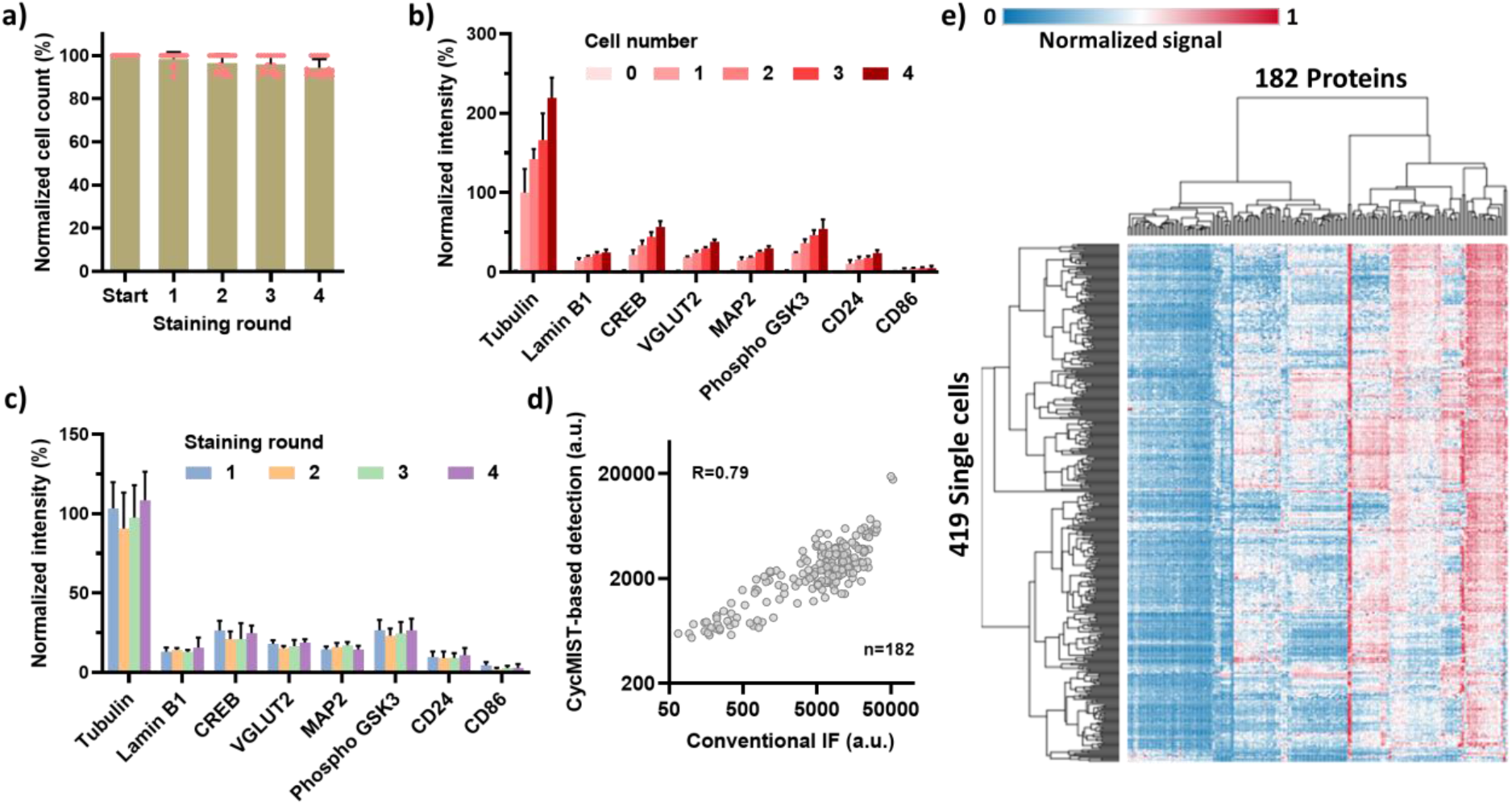
Single-cell CycMIST assay on N2a cell line. a) Quantification of cell loss through 4 rounds of the assay on the same cells in the PDMS microwells. Error bars represent 25 times of repeats. b) Assay sensitivity for increased number of cells loading in microwells. Error bars are standard deviations of signals from 10 independent microwells. c) Reproducibility of CycMIST analysis of 8 proteins for 4 staining rounds on the same single cells. Error bars are standard deviations of signals from 15 single cells. d) Correlation of 182 protein expression levels between the single-cell averages measured by CycMIST and the population levels measured by conventional immunofluorescence staining methods. The CycMIST-based signal is obtained by averaging the signal of 100 single cells. The correlation coefficient R equals 0.79 for the two methods. e) Heatmap of unsupervised clustering result of a single-cell CycMIST assay using a whole panel of 182 proteins with 4 staining rounds. Both rows and columns are clustered by Euclidean distance and complete linkage.

The validated platform has been applied to measure 182 proteins in single neurons. This large panel includes proteins for typical phenotype markers of major brain cell types, AD hallmark pathway, neuron functions, enzymes, canonical signaling pathways (e.g., insulin signaling, calcium signaling, JAK/STAT signaling, autophagy signaling and apoptosis), and transcriptional factors. They were selected from Mouse Brain of Allen Brain Atlas, AlzPathway and publications related with 5xFAD mouse model (Table S2)^38, 39, 40^. All the antibodies targeting those proteins were conjugated with their corresponding cDNAs, and the whole set was divided into 4 panels for 4 staining rounds of CycMIST experiments (Table S1). The antibodies that can potentially interfere with each other (i.e., affinity to the protein in the phosphorylated vs. unphosphorylated state) were intentionally separated in different panels. The averaged single-cell result for each protein is compared with that by conventional bulk assay through immunofluorescence staining (Figure 3d). The results are relatively comparable to each other, despite that they are based on two different assays. Both cells and proteins from the single-cell assay was clustered into groups (Figure 3e). It is found that the surface markers are enriched on the left of the heatmap, while enzymes and essential proteins to cells are highly expressed on the right. This is expected because the cell line N2a does not express a broad spectrum of brain cell markers while enzymes and cytoskeleton relevant proteins are highly abundant in typical animal cells. Autophagy related 5 (ATG5), HMG synthase 1 (HMGCS1) and HMG-CoA reductase (HMGCR) are apparently the highest detectable proteins for the N2a cells in this study.

### Single-cell functional proteomics study of mouse brain cells

The single-cell CycMIST assay has been used to analyze primary mouse cells from wild-type mice (WT) and 5XFAD transgenic mice (AD). The 5XFAD mouse model is a transgenic mouse that overexpresses the human amyloid precursor protein (APP) and human presenilin-1 containing a total of 5 familial AD mutations, and amyloid accumulation is observable at 2 months of age^41^. Neuron loss and memory deficits already occur at 6 months of age ^41^. In this report, the pre-frontal cortices of the 4.5-month-old mice were surgically dissected and dissociated into single-cell suspensions containing all brain cell types, and they were immediately loaded and fixed within the PDMS microwell. The same 182 protein panel was used to analyze the cells to collect 721 AD single-cell data and 675 WT single-cell data (Figure 4a).

**Figure 4.**
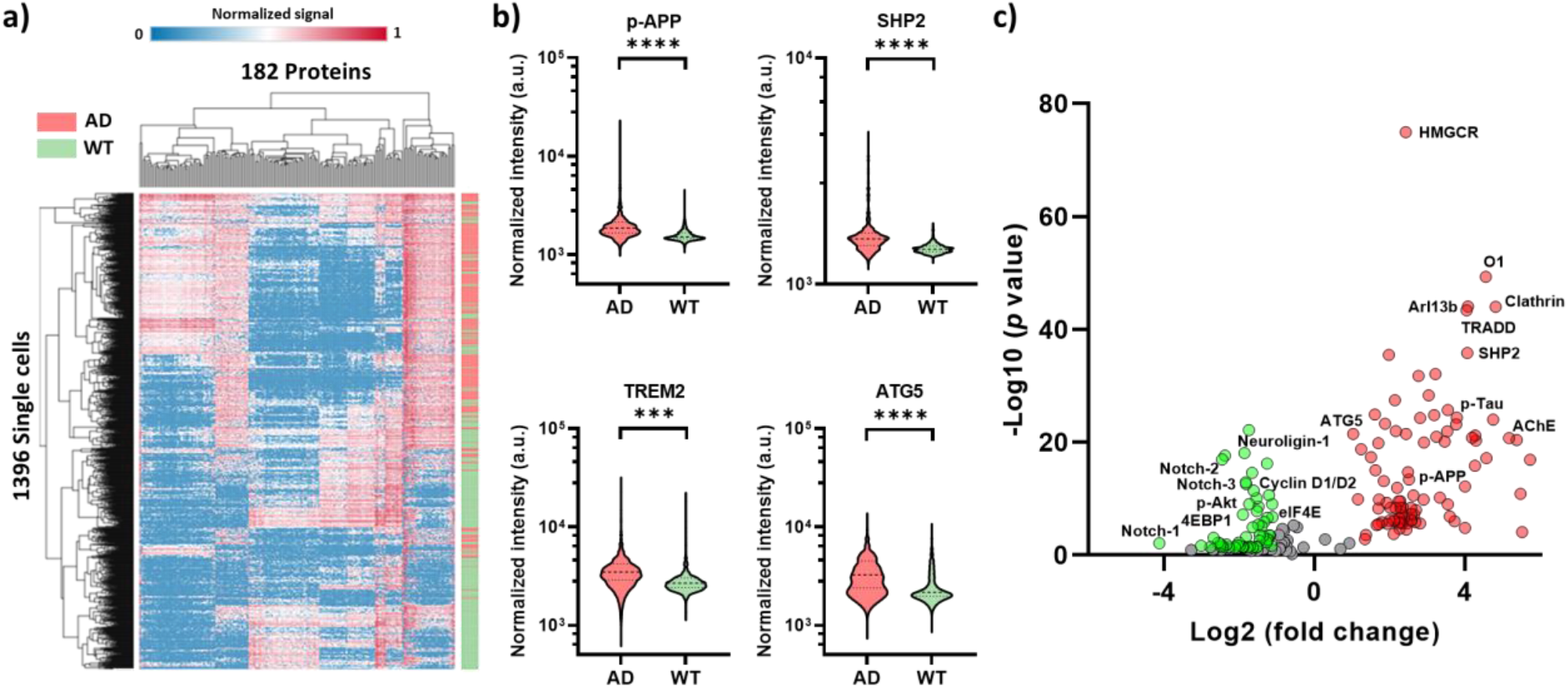
Comparison of AD and WT cortical cells through the single-cell CycMIST assay. a) Heatmap of unsupervised clustering for both AD and WT cells. Color bar on the right indicates the distribution of the single AD cells (red) and single WT cells (green). The single-cell data above background + 3 S.D. are used for clustering by Euclidean distance and complete linkage methods. b) Violin plots of 4 representative proteins related with AD development and comparison between AD and WT cells. ***: *p*-value < 0.001; ****: *p*-value < 0.0001. c) Volcano plot of differentially expressed proteins in AD and WT cells. Red dots represent proteins expressed at high levels in AD cells, and green dots are proteins expressed at high levels in WT cells. Y axis is − log10 (*p*-values) while X axis shows log2 fold change values.

*p*-APP levels are significantly higher in AD brain cells than WT cells (Figure 4b and Figure S8), which is consistent with the nature of the 5XFAD mouse model. The neurodegeneration and neuroinflammation related proteins including Src homology 2 (SH2) domain–containing phosphatase 2 (SHP2), Triggering receptor expressed on myeloid cells 2 (TREM2), and ATG5 are also expressed at a higher level in AD cells, compared to WT cells^42, 43, 44^. To visualize the protein expression profiles of single cells and compare the two samples, background noise was excluded and only the real signal of detection was subject to clustering analysis (Figure 4a). The single-cell profiles of AD and WT cells tend to be enriched in certain sub-clusters of proteins respectively, while in some sub-clusters they are indistinguishable. To identify the individual proteins that are differentially expressed by the cells, we took advantage of large sample sizes (hundreds of cells) to obtain the fold change from WT to AD for every protein with a *p* value (Figure 4c). In general, the AD cells have more detectable proteins than the WT cells. This might be attributed to an upregulation in cellular responses to neuronal death as well as subsequent glial activation.

## Discussion

The CycMIST assay is highly quantitative by its nature and thereby can estimate the number of proteins in single cells. Since a conjugate is labeled with ~3 cDNAs in average and each microwell is 225 pL, the estimated LOD for protein detection is 180 protein copies per cell, which is comparable to other multiplexed single-cell technologies^23, 25, 29, 33^. The high sensitivity is attributed to the high-density DNA on the MIST array and the small size of the microwells. However, this estimation must assume most of proteins are accessible for antibody binding, and the protein and the antibody are 1:1 ratio. For that reason, we have not converted the single-cell detection signals to protein quantities yet.

The 182 proteins selected in the assay are highly diversified: they are distributed in various compartments of a cell with varied functions. The single-cell assay results show high heterogeneity in the expression of those proteins for both N2a cell line and mouse primary cells. Just like other typical single-cell omics technologies, the tissue needs to be digested to isolate cells before the analysis. Although an optimal protocol was used to separate neurons from mouse brains, most likely a good portion of the axon and dendrites were lost and only soma was analyzed. This is still better than typical single-cell transcriptomics techniques that isolate the nuclei for sequencing. Further improvement will lie on modification of the tissue dissociation protocol. Nevertheless, we were still able to capture certain upregulated proteins in cells dissociated from AD mouse tissue, which is in line with what is known in literature about pathways upregulated in the AD brain. The typical proteins elevated in AD pathology are statistically higher expressed in our result, like p-APP and certain glial activation markers, while proteins related to cell growth and division signaling are suppressed. Through gene ontology analysis (Table S3 and S4), it is found the ROCKs signaling and chylomicron clearance are significant activated in AD cells. ROCK protein levels in neurons increase together with the activated amyloid β in mild cognitive impairment of AD brains^45^. Chylomicron contains apolipoprotein B (apoB) that may be involved in encapsulation of amyloid β and lipid metabolism in early onset of AD^46, 47^. In addition, the analysis result shows MAPK pathway receives high negative feedback signaling that is related with Frs2 mediation and PI3K regulation. Thus, the MAPK pathway might be constantly activated in our AD cells. This is consistent with the reports that MAPK pathway is frequently altered in AD development and play a key role in neuroinflammation^48^. Our assay may provide further insights of the mechanisms in MAPK participation and potential targets for treatment.

In summary, we demonstrate the single-cell CycMIST technology as a functional proteome profiling tool. It breaks the bottleneck of limited multiplexity in single-cell protein measurement where only dozens of proteins are detected. Single-cell CycMIST can assay up to 50 proteins every 2.5 h and ~200 proteins in 1 day, which is much faster than other similar types of technologies. Higher multiplexity should be achievable simply by increasing the round number, since even 20 rounds of immunofluorescence staining of cells can still produce uncompromised staining result^16, 18^, and for CycMIST, each round is a separate assay. Further development of the CycMIST technology will be focused on automation and data acquisition. Our microchip is amenable to integration with an instrument that can handle liquids and scan the entire array. Development of advanced data acquisition will be needed to standardize the process to be like flow cytometry, which has the hardware gating system and associated software to improve data quality and lower detection noise. The single-cell CycMIST technology will potentially become a routine functional proteome tool broadly applied in various biomedical fields for mechanistic studies and diagnostics.

## Methods

### Fabrication of PDMS microwell chip

Fabrication of polydimethylsiloxane (PDMS) chips followed the conventional methods in soft lithography^49^. Briefly, a chrome photomask (Front Range Photomask) was used to pattern a layer of features with 40 μm thickness on a 4” silicon wafer (University Wafer) by photolithography and photoresist SU-8 2025 (Kayaku Advanced Materials). The resultant mold was then pretreated with trimethylchlorosilane (TMCS; Sigma Aldrich) for 30 mins to facilitate PDMS separation. Afterward, a mixture of PDMS prepolymer and curing agent (Salgard 184; Dow Corning) with a ratio of 10:1 was cast on the mold. Air bubbles were removed via vacuum desiccator for 1 h, and the PDMS mixture was baked in an oven at 80°C for 2 h. The cured PDMS elastomer was peeled off and cut into the appropriate size for further use. Each array on the PDMS chip contains thousands of microwells for cell loading, and each microwell features with a dimension of 75 μm (length) × 75 μm (width) × 40 μm (depth).

### ssDNA modification on microbeads

Polystyrene microbeads (2 μm; Life Technologies) with amine group were coated with poly-L-lysine (PLL; Ted Pella) first to amplify functional groups on the beads surface. Specifically, 100 μL of 2% amine bearing microbeads was treated with 100 μL of 10 mM bis(sulfosuccinimidyl)suberate (BS3; Pierce) crosslinker solution for 20 mins. Subsequently, the microbeads were washed with Milli-Q water and then mixed with 1 mL of PLL in phosphate buffered saline (PBS; pH 8.5) buffer overnight on a vortex shaker. The PLL-funtionalizaed microbeads were rinsed with PBS buffer and then reacted with a mixture of amine-ended oligo ssDNA (30 μL; 300 μM; Integrated DNA Technologies), BS3 (130 μL; 2 mM), DMSO (69 μL) and PBS (31 μL) buffer solutions for 4 h. Finally, the microbeads were throughly washed with Milli-Q water and resupended to the orignal concentration for further use.

### Patterning monolayer of the MIST array and characterization

All the 50 ssDNA-functionalized microbeads were mixed in equal portion first (14 μL for each) and then mixed with 300 μL of the blank microbeads to reduce the signal overcrowding during imaging. After sonication for 10 mins, 10 μL of mixed microbeads solution was pipetted onto a 0.5 cm × 0.5 cm surface area of PDMS slab and left to dry. The dried microbeads were carefully transferred onto a cleanroom adhesive tape (VWR) which was attached to a plain glass side. Afterwards, the array was sonicated for 5 mins to remove the excess layers of microbeads, leading to the formation of the uniform monolayered MIST array. The MIST arrays were prepared in batches of 40 to 50, which were then stored dry at 4°C for later use.

The MIST array was characterized by using a cocktail of Cy5-labelled cDNAs solutions (200 nM in 3% BSA/PBS), which was applied to the array and then incubated for 1 h, followed by PBS washing and imaging under a fluorescence microscope. The intensities of each type of microbeads were measured by ImageJ. The total number of the ssDNA-microbeads and the number of each type of ssDNA-microbeads for each 75 μm × 75 μm array were quantified by a MATLAB program developed in the lab. To make calibration curve for each type of ssDNA-microbeads, various concentrations of Cy5 tagged cDNA (1 pM, 10 pM, 100 pM, 1 nM and 10 nM) were added onto the MIST array, and their fluorescence intensities were quantified. A logistic fitting function was applied to each calibration curve after taking log scale of the cDNA concentrations. The limit of detection (LOD) for the 50 types of ssDNA-microbeads are calculated by extrapolating the background fluorescence intensity plus three times of its standard deviations using the fitting curve.

### Crosstalk validation of oligo DNAs

DNA sequences were designed by a Python program that sets Tm at least 50°C, >53% GC content, <4 internal dimers and Tm <5°C of cross reactivity. Over 150 candidate oligo DNAs were selected and experimentally screened to narrow down to a panel of 50 oligo DNAs. The cross-reactivities of the 50 oligo DNAs were validated by using fluorophore labelled cDNAs. Typically, to investigate the crosstalk between Seq *n* (*n* means the n type of DNA) with other 49 DNAs, a MIST array only containing Seq *n*-modified microbeads and blank microbeads was fabricated according to the aforementioned process. A Cy5-labelled Seq *n*’ cDNA solution (200 nM in 3% BSA/PBS) was added onto the array and incubated for 1 h. The array was then thoroughly washed with PBS and imaged by a fluorescence microscope, which was denoted as positive signal. On the other hand, a 200 nM cocktail solution of the other 49 Cy5-labelled cDNAs (expect for Seq *n*’) was applied to the array, followed by the same procedure above, and the obtained picture was denoted as crosstalk signal. The fluorescence intensities of the positive and crosstalk signals were quantitatively analyzed by ImageJ.

### Preparation of cDNA-Dye conjugates

Three fluorescent colors (Alexa Fluor 488, Cy3 and Cy5) of cDNA-dye conjugates were used for the decoding of 50 different ssDNA-microbeads. The Cy3- and Cy5-labelled cDNAs were directly purchased from Integrated DNA Technologies, and the Alexa Fluor 488-labelled cDNAs were synthesized in the lab. Briefly, 50 μL of 200 μM cDNAs were incubated at pH 8.5 with 50 equivalents of Alexa Fluor 488 NHS Ester (Life Technologies) in 30% DMF for 4 h. Afterwards, excess chemical reagents were removed by Zeba spin desalting column (7K MWCO; Thermofisher), with PBS as a wash buffer. The Alexa Fluor 488-labelled cDNAs were diluted to 10 μM and stored at −20 °C for later use.

### Preparation and purification of UV-cleavable, biotinylated cDNA-antibody conjugates

All the antibodies (BSA-free) details and their corresponding cDNA barcodes used in this study were summarized in Table S1 and Table S2 in the supplementary information. Prior to cDNA conjugation, all antibodies were tested on N2a cells by conventional immunofluorescence microscopy and then compared with the information provided by vendors and literature to ensure the binding and localization of each antibody with respect to its target. Custom designed oligo DNAs were purchased from Integrated DNA Technologies and used as received.

The conjugation of biotinylated cDNA with antibody was achieved through click chemistry reaction. Reactions should be covered in foil or in a dark room due to the light sensitivity of UV-cleavable crosslinkers. In detail, antibodies were first concentrated to 1 mg/mL or higher using Amicon Ultra centrifugal filter (10 K MWCO; EMD) to improve the conjugation efficiency. Both antibodies and cDNAs were then solvent exchanged to a PBS buffer with pH 8.5 by Zeba spin desalting column (7K MWCO; Thermofisher) to remove sodium azide and reach the optimal pH for efficient amine labelling. Afterwards, 50 μL of 1 mg/mL antibodies were reacted with 1 μL of 10 mM UV-cleavable azido-NHS ester (30 equivalents to antibodies; Click Chemistry Tools) for 2 h, while 30 μL of 200 μM cDNAs were incubated with 1.2 μL of 100 mM UV-cleavable DBCO-NHS ester (20 equivalents to cDNAs; Click Chemistry Tools) in 30% DMF for 4 h. The azido-antibodies and DBCO-cDNAs were buffered exchanged to pH 7.4 PBS using Zeba spin desalting column (7K MWCO) to remove excess regents. The DBCO-cDNAs were mixed with their corresponding azido-antibodies for reaction overnight at 4°C, resulting in the formation of cDNA-antibody conjugates. The conjugates were purified using a FPLC workstation with Superdex® 200 gel filtration column at 0.5 min/min isocratic flow of PBS buffer. Only a portion of the conjugate peak is selected to ensure high cDNA loading to each antibody. Finally, the desired products were collected from FPLC elutions and then concentrated to 0.3-0.5 mg/mL and stored in 4°C for further use. The absorbance spectrum of UV-cleavable biotinylated cDNA-antibody conjugates were measured by a Nanodrop spectrophotometer (Thermofisher) to determine the degree of labeling (DOL) by following equation^32^:

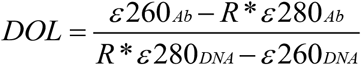

Where ε260_Ab/DNA_ and ε280_Ab/DNA_ are the extinction coefficients of each cDNAs and antibodies at both 260 nm and 280 nm, respectively. R is the absorbance ratio (260/280 nm) of the cDNA-antibody conjugates.

### Cell preparation and immunofluorescence staining

Mouse neuroblastoma Neuro-2a (N2a) cells (ATCC) were cultured in Dulbecco’s modified Eagle’s medium (DMEM; Life Technologies) containing 10% (v/v) fetal bovine serum (FBS), 100 μg/ml streptomycin sulfate, and 100 U/ml penicillin G sodium at 37°C in a humidified atmosphere with 5% CO_2_. N2a cells were passaged twice a week with a cell density of ~80-90%. For the induction of neuronal differentiation, N2a cells were plated at a density of 2 × 10^4^/cm^2^ and maintained in the standard growth medium for 24 h. Next day, the N2a cells were stimulated by a reduced serum medium (DMEM supplemented with 2% FBS) in the presence of 20 μM Retinoic acid (RA; Sigma-Aldrich). The medium was refreshed every 24 h. The cells having one or more neurites of a length more than twice the diameter of the cells body were defined as differentiated.

Prior to cell staining, all the buffers and solutions were first warmed up to room temperature. Then the cells were rinsed with PBS buffer, fixed with 4% formaldehyde in PBS for 20 mins, permeabilized with 0.1% Triton-X100/PBS for 7 mins and washed three times with PBS buffer. For the conventional immunostaining by the unmodified antibodies, the fixed cells were incubated with 5% goat serum (Cell signaling) in PBS for 1 h, stained with 10 μg/mL of primary antibodies in 5% goat serum for 1 h, incubated with fluorophore-conjugated secondary antibodies in 5% goat serum for 1 h and then stained with 5 μg/mL of Hoechst 33342 (Pierce) in PBS for 15 mins. The cells were washed three times with PBS buffer between the steps. For staining by the cDNA-antibody conjugates, the fixed cells were incubated with a blocking buffer (10% goat serum, 2% BSA, 1 mg/mL Salmon Sperm DNA and 0.1 % Tween 20 in PBS) for 0.5 h to minimize non-specific antibody and DNA binding. After washing three times with PBS, the cells were stained with 10 μg/mL of cDNA-antibody conjugates in the blocking buffer for 1 h, followed by incubation with fluorophore-conjugated secondary antibodies for 1 h and then stained with 5 μg/mL of Hoechst 33343 in PBS for 15 mins. Finally, the cells were imaged under a fluorescence microscope.

### UV cleavage and cyclic immunostaining

Eight different UV-cleavable Cy5 labelled cDNA-antibody conjugates (anti-tubulin/Lamin B1/CREB/VGLUT2/MAP2/p-GSK3/CD24/CD86) were used to label cells according to the immunostaining procedure above. For kinetic measurements, the plates with cells were exposed to a UV light (365 nm; Thorlabs CS2010 UV Curing LED System) for different durations and then washed thoroughly with PBS buffer. Fluorescence images of the cells before and after UV irradiation under different exposure time were taken using a fluorescence microscope, and the fluorescence intensities of each image were quantitatively measured by ImageJ.

Cyclic immunostaining was performed in a parallel and sequential manner in 96-well plates. Briefly, for each of the eight conjugates, each staining round consisted of four steps: 1) blocking; 2) staining; 3) imaging; 4) washing. The first three steps were identical to the staining procedure as described above. The washing step was achieved by incubating the stained cells with 200 μL of regeneration buffer (composed of 0.5 M L-Glycine, 3 M Urea, 3 M Guanidinum chloride in Milli-Q water, pH 2.8) three times for 2 mins each^16^. Afterwards, the cells were stained with fluorophore-conjugated secondary antibodies to verify that the conjugates were completely eluted from the cells. Following the washing step, the cells were washed with PBS and incubated with the blocking buffer for 1 h to proceed the next staining round. This staining round was repeated four times to complete the cyclic immunostaining. Fluorescence images of the cells after each staining and washing step of the four cyclic staining rounds were recorded and quantitatively analyzed.

### Multiplexed single-cell protein detection and decoding process

The PDMS microwell were first treated with a plasma cleaner (Harrick Plasma) for 1 min and then incubated with 0.5% Pluronic® F-127 (Sigma Aldrich) in Milli-Q water for 30 mins to reduce nonspecific conjugate adsorption on the surface. After cleaning with PBS, a cell suspension containing ~10,000 cells in culture medium was applied onto the PDMS surface, and the PDMS slab was then placed on a shaking table with a speed of 40 rpm/min for cell loading. The excess cells that were not attached to the microwell surface were gently removed by pipetting. The cells inside of the microwells were stained with a cocktail of UV-cleavable biotinylated cDNA-antibody conjugates according to the immunostaining protocol as described above. After the completion of the staining, the chip was washed three times with PBST containing 0.1% Tween 20 and 5% goat serum. A glass slide carrying a MIST array was carefully mated with the PDMS chip on a clamp to completely isolate the cells in the microwells. The whole device was exposed to a UV light for 15 mins to release the biotinylated cDNAs and then left for another 1 h to allow the hybridization of cDNAs with ssDNA on the MIST array. The MIST array was then separated from the PDMS microwells and washed with 3% BSA in PBS. 150 μL of 10 μg/mL streptavidin-Alexa Fluor 647 dye (Life Technologies) in 3% BSA/PBS solution was added on the MIST array for signal visualization, which was corresponding to protein signal. Meanwhile, the cells in the PDMS microwells were washed with the regeneration buffer for the next staining round by another different cocktail of conjugates.

The fluorescence images of the protein signal on the MIST array were recorded and then proceeded to the decoding process. The decoding procedure consisted of four cycles using three different colors of cDNA-dyes, which can identify maximum 3^4^ (81) types of microbeads/proteins. For the decoding of the first cycle, 150 μL of 2 M NaOH solution was applied onto the array to dissociate double stranded DNAs for 1 min. After washing with saline-sodium citrate buffer (SSC) for three times, 150 μL of a cocktail of cDNA-dye conjugates (Cocktail I, 200 nM) in hybridization buffer (40% formamide and 10% dextran sulfate in SSC buffer) was added on the array and incubated for 1 h. The array was washed with SSC buffer and subsequently imaged under a fluorescence microscope, which was denoted as Cycle 1 signal. The procedures of the second, third and fourth decoding cycles were similar to that of the first cycle, except that Cocktail II, Cocktail III and Cocktail IV of cDNA-dye conjugates were used to label the array, resulting in the decoding signals of Cycle 2, Cycle 3 and Cycle 4, respectively. All the images of the Cycle 1 signal to Cycle 4 signal were registered together to determine the order of fluorescent color change of each microbead by a Matlab program developed in the lab.

### Mouse brain sample preparation

All procedures were conducted in accordance with the US National Institutes of Health Guide for the Care and Use of Laboratory Animals and were approved by the Institutional Animal Care and Use Committee (IACUC) at Stony Brook University. Brains of male wild-type (WT) and C57BL/6 (5XFAD) transgenic mice with ages of 4.5-month-old were used in this sudy. The cortex regions were then dissected and placed on ice in bucket for immediate use. A commercial Adult Brain Dissociation Kit (Miltenyi Biotec) was used to dissociate the cortex samples into individual cells by following the separation protocol from the manufacturer. The dissociate cells were stained by a 2 μM of Calcein AM dye (Life Technologies) solution to identify the cell viability and then loaded into PDMS microwell chips for the multiplexed single-cell protein detection analysis.

### Image quantification and registration

A Nikon Ti2 inverted fluorescence microscope equipped with a motorized stage was used to automatically take images for the entire PDMS microwells and the MIST array. Bright field and phase contrast images were utilized to identify the microwell address and microbead locations. For the fluorescence images on the Hoechst, Alexa Fluor 488, Cy3 and Cy5/Alexa Fluor 647 channels, the light were filtered with beam splitters and emission filters controlled by a wavelength switcher. All the images were saved as the format of 16-bit ND2 in the Nikon software (NIS-Elements AR Microscope Imaging Software) and then were passed to MATLAM program for further registration and processing.

An in-house MATLAB program code was utilized for image overlapping, registration and signal analysis. The digitalized images for each dataset on a 75 μm × 75 μm array composed of protein signal images, bright-field images and three different color images from the four decoding cycles. Bright images from protein signal were read first as a reference to generate registration information, and the same registration information were used to align images from the protein signal and four decoding cycles. All the aligned images were then compiled into multi-image stacks to identify the order of fluorescent color change for each microbead and quantify the fluorescence intensity of individual microbead on the protein signal. The protein ID of each microbead was determined by comparing the color order with the proteins in the decoding design (Table S1). The microbeads detecting the same protein were grouped and their fluorescence intensities were averaged for each 75 μm × 75 μm array. After compiling the information of cell number counting on each microwell or microbead array, a final dataset was produced including the results of cell number (zero or one cell) and the proteins detection signal and their corresponding fluorescence intensities.

### Data analysis and statistics

Unless otherwise stated, all measurements were performed at least three duplicates (n≥3) and the data were presented as mean ± S.D (standard deviation). For single cells analysis, the detected protein signal in the microwells of zero cell was taken as the background signal. The mean intensity plus 3 × standard deviation of the background signals was considered as the confident threshold, and the proteins signals of each microbead array above the threshold were regarded as the true signals of the detected proteins. Unpaired, 2-tailed T-test was used to evaluate the statistical significances with GraphPad Prism 8.0, which were considered at the *p*-value < 0.001 (***) and *p*-value < 0.0001 (****) levels. The same software was used to generate violin plots. For plotting the 182 panel, a python program with seaborn library was developed to generate the figures. The univariate analysis of protein intensities for AD and WT samples was based on fold-change values and the threshold of significance by building a volcano plot (setting: p-values <0.05, fold change >2.0). Heatmaps and unsupervised clustering were generated with the Morpheus program (https://software.broadinstitute.org/morpheus).

## Data availability

All the data supporting the study are available within the article and its Supplementary Information files, and from the corresponding author upon reasonable request.

## Acknowledgements

This work was supported by the National Institutes of Health R21AG072076 (JW), R01GM128984 (JW), R01NS096275 (YML), RF1AG057593 (YML), and the JPB Foundation (YML). Authors also acknowledge the MSK Cancer Center Support Grant/Core Grant (Grant P30 CA008748).

## Author contributions

J. W and L. Y designed the study, performed the experiments, provided data analysis and wrote the manuscript. T.J and Y.M. L provide mouse samples and experimental design. A. B. optimized the code of MATLAB program. J. L assisted in the data processing for the mouse brain samples. All the authors reviewed the manuscript and approved the submission.

