## Supplementary information for "Cyclic Microchip Assay for Measurement of Hundreds of Functional Proteins in Single Neurons"

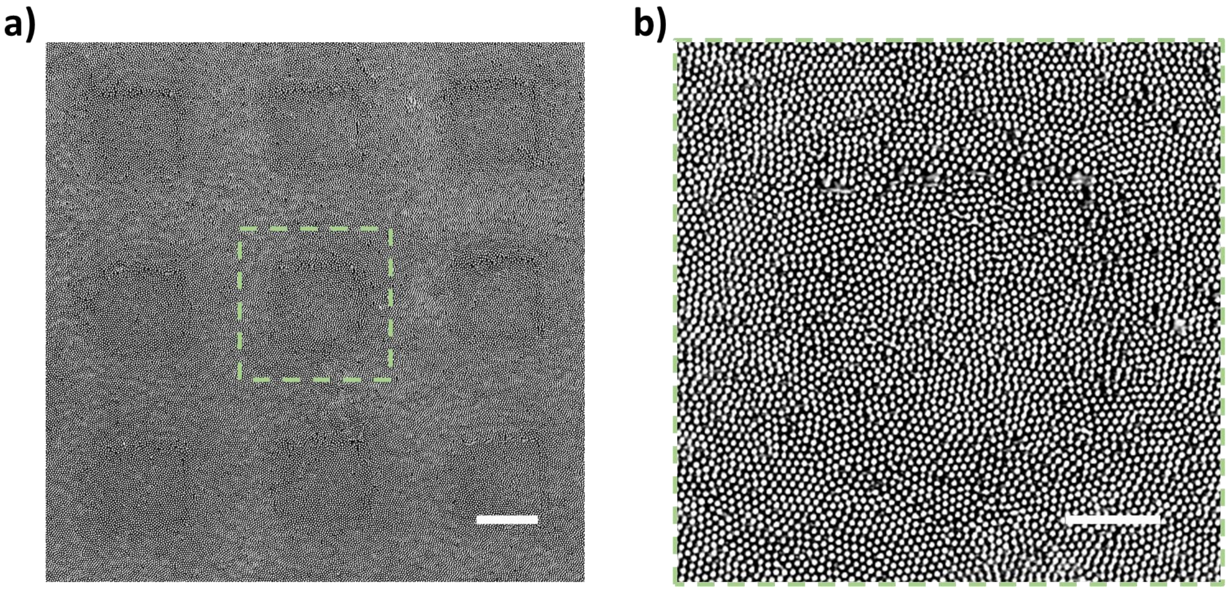

**Figure S1.** a) Bright field images of a large-scale microbead array after clamping with a PDMS microwell chip. Scale bar = 50 μm. b) Zoom-in images of the green dotted frame in a. Scale bar = 20 μm.

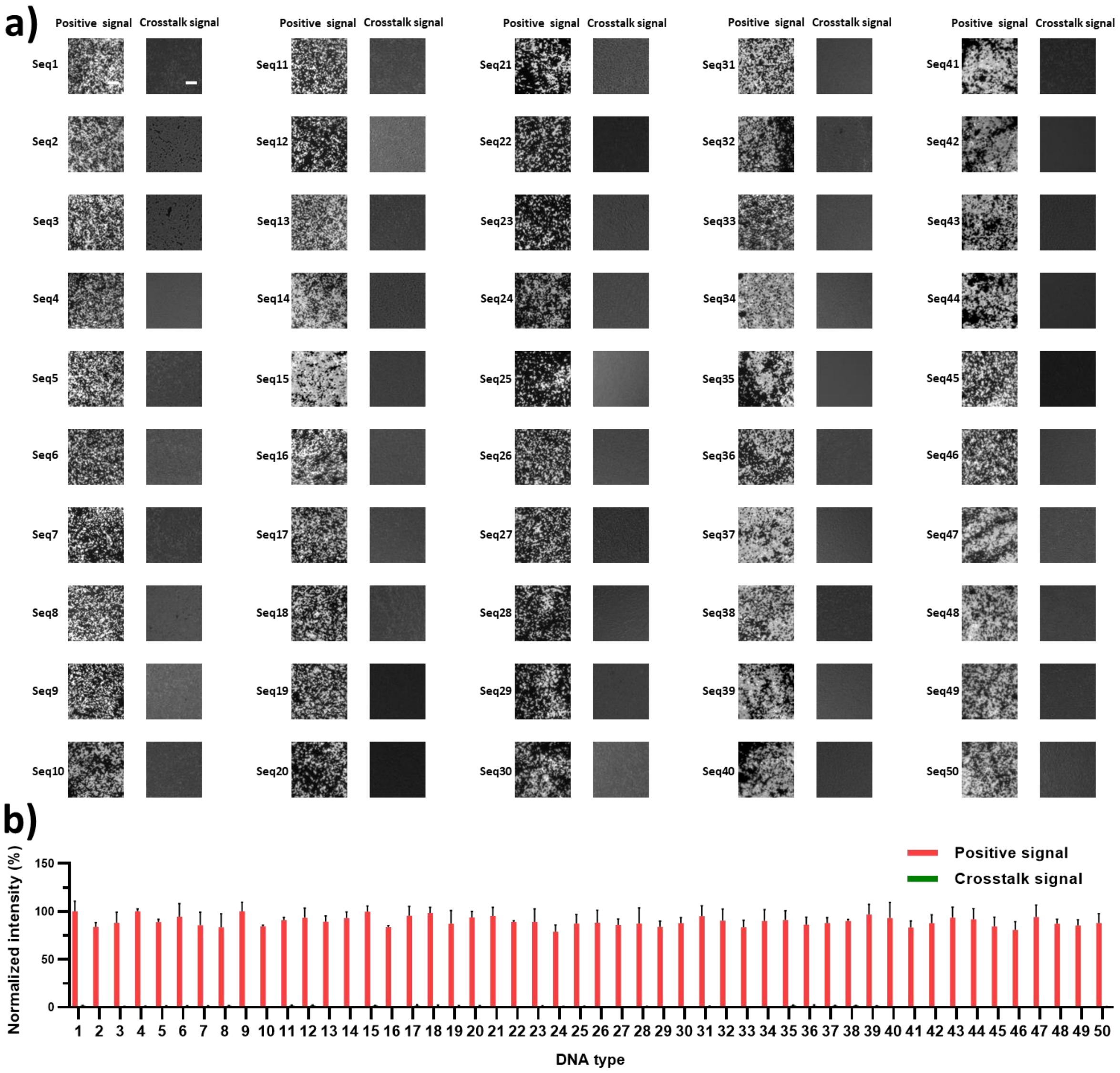


**Figure S2.** Crosstalk validation of 50 different types of oligo DNAs. a) Fluorescence images of positive signal and crosstalk signal for each tested DNA. b) The fluorescence intensity of each images in a, which were analyzed quantitatively using ImageJ and normalized by the intensity of positive signal for Seq1 DNA. Data are presented as mean ± SD, n = 3. Error bars are within symbol size if not shown.

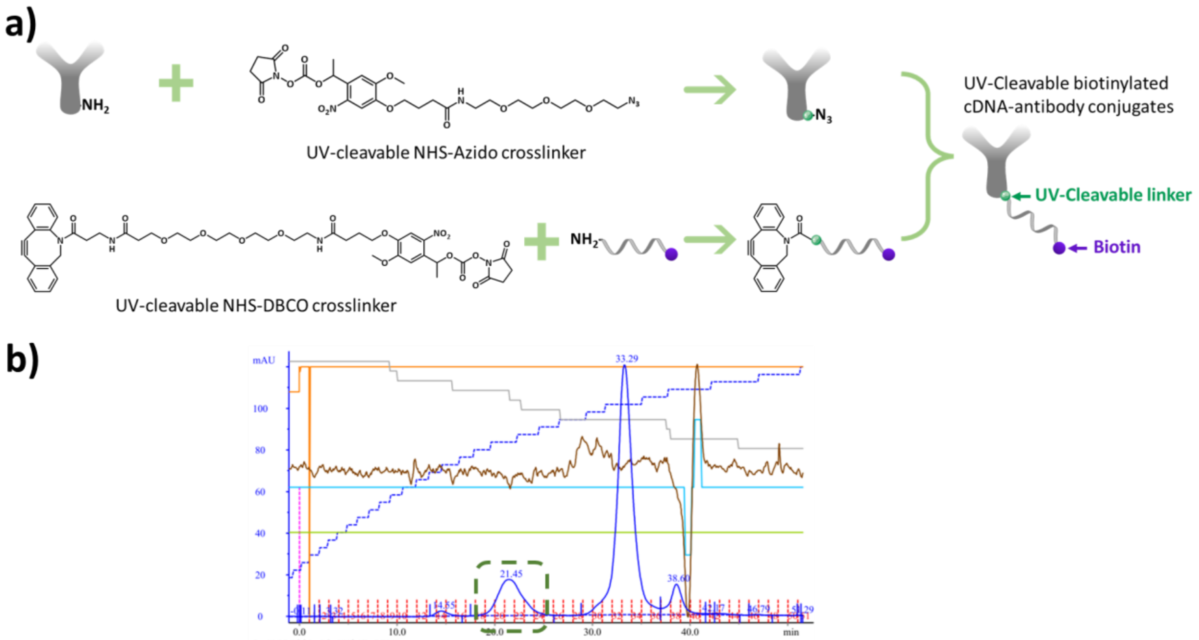


**Figure S3.** a) Synthetic scheme for the preparation of UV-cleavable biotinylated cDNA-antibody conjugates. b) Typical FPLC purification curve of the conjugates, the green dotted frame indicates the peak of the purified conjugate, where the location in the chromatography is correlated with the number of chemical loads.

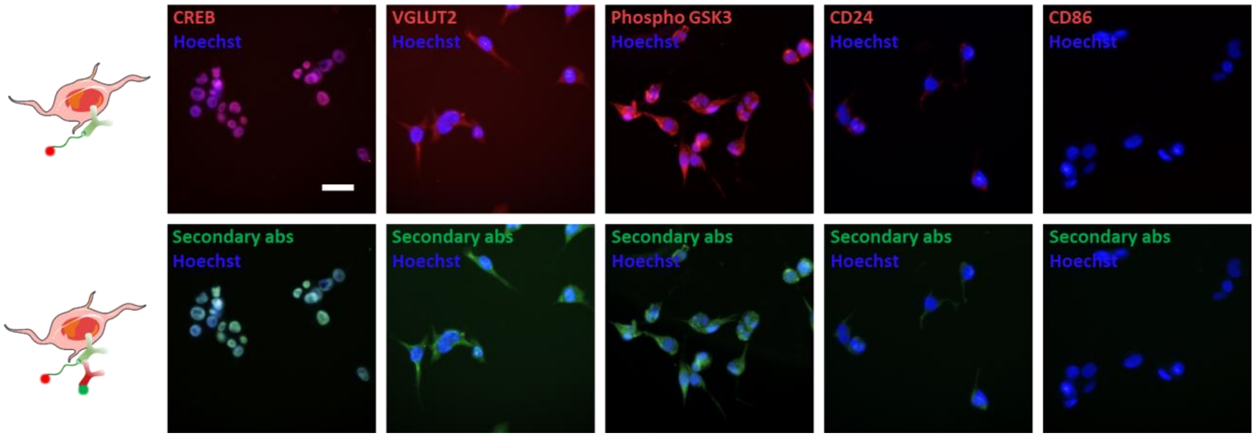


**Figure S4.** Validation of staining specificity of conjugates. Up panel: fluorescence images of the differential N2a cells after incubation with cy5-labeled cDNA conjugated to anti-CREB/VGLUT2/Phospho-GSK3/CD24/CD86 antibodies. Bottom panel: fluorescence images of the stained cells after incubation with Alexa Fluor 488-labelled secondary antibodies. Scale bar = 20 μm.

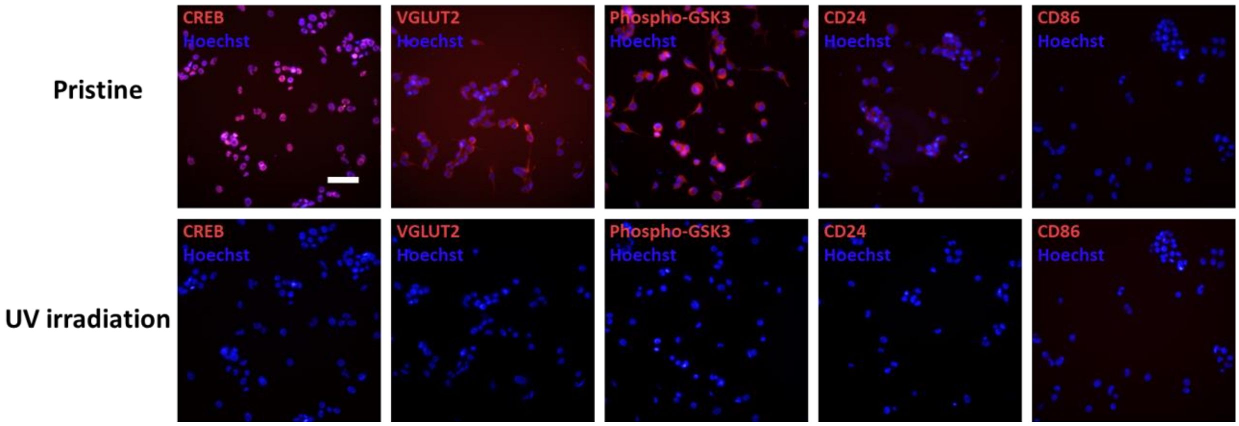


**Figure S5.** UV-cleavage efficiency of ssDNA-antibody bonds on conjugates. Up panel: fluorescence images of the differential N2a cells after incubation with anti-CREB/VGLUT2/Phospho-GSK3/CD24/CD86 antibodies that were tagged with cDNA-Cy5. Bottom panel: fluorescence images of the stained cells after 15 min of UV-light irradiation, demonstrating that cDNA barcodes were released from the stained cells. Scale bar = 50 μm.

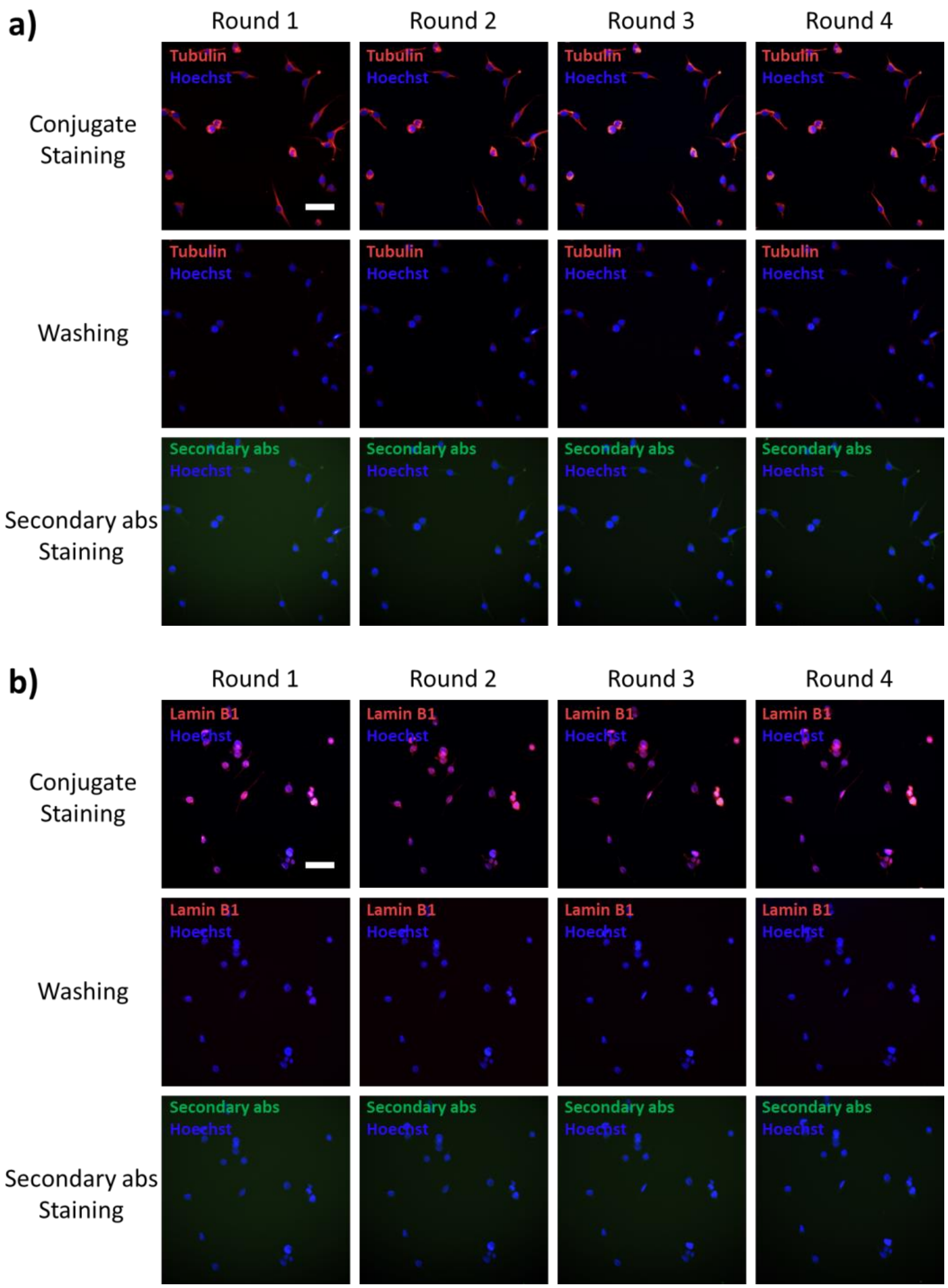

a)
Round 1
Round 2
Round 3
Round 4
Conjugate Staining
Washing
Secondary abs Staining
Tubulin
Hoechst
Secondary abs
b)
Round 1
Round 2
Round 3
Round 4
Conjugate Staining
Washing
Secondary abs Staining
Lamin B1
Hoechst
Secondary abs

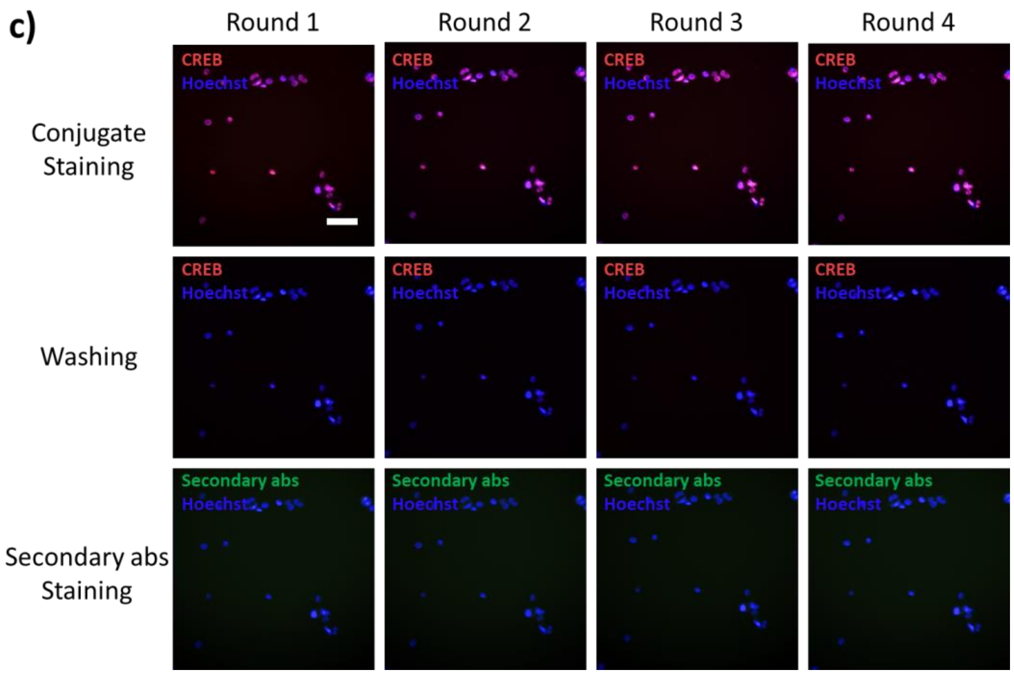
c)
Round 1
Round 2
Round 3
Round 4
Conjugate Staining
Washing
Secondary abs Staining
CREB
Hoechst
Secondary abs

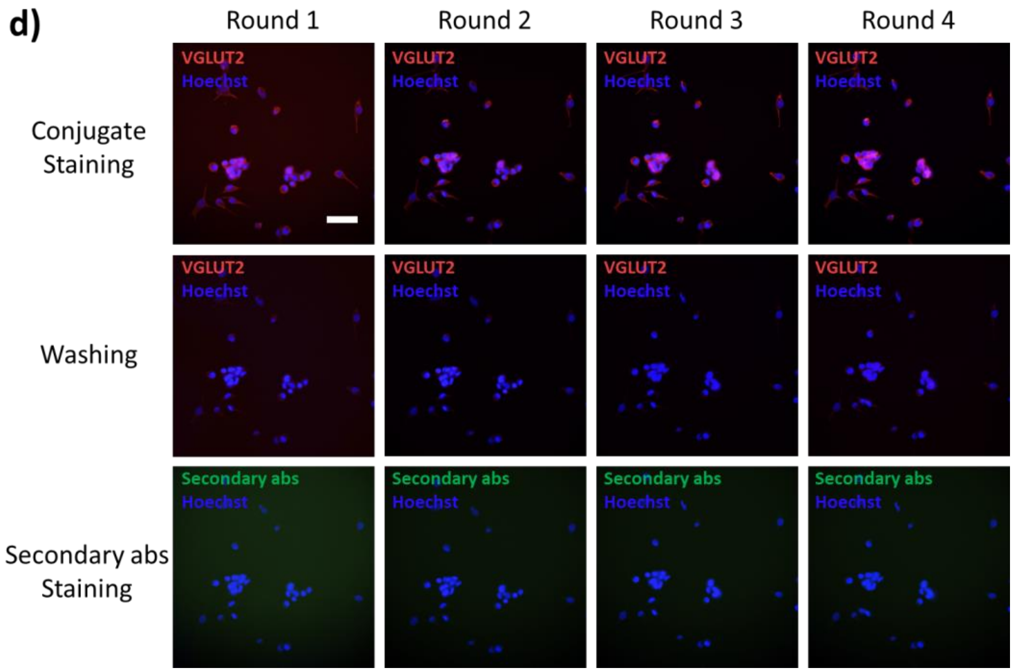
d)
Round 1
Round 2
Round 3
Round 4
Conjugate Staining
Washing
Secondary abs Staining
VGLUT2
Hoechst
Secondary abs

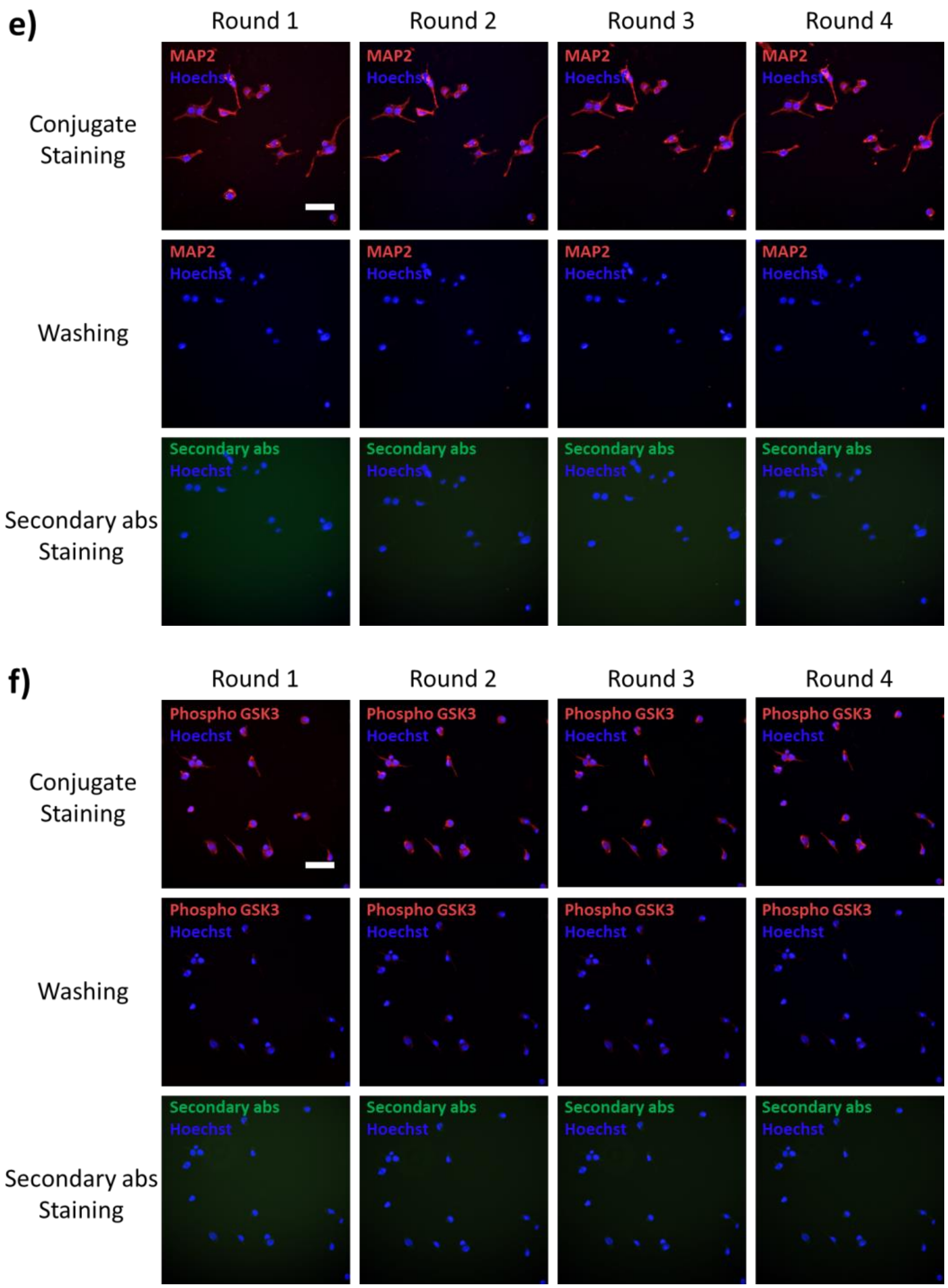
e)
Round 1
Round 2
Round 3
Round 4
Conjugate Staining
Washing
Secondary abs Staining
MAP2
Hoechst
Secondary abs
f)
Phospho GSK3

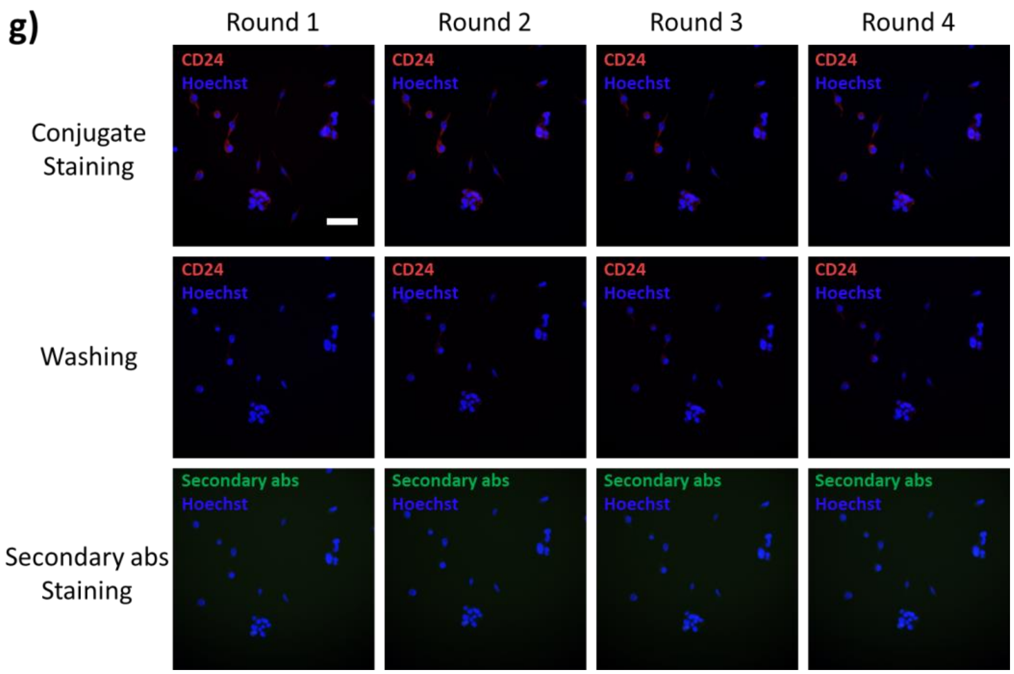


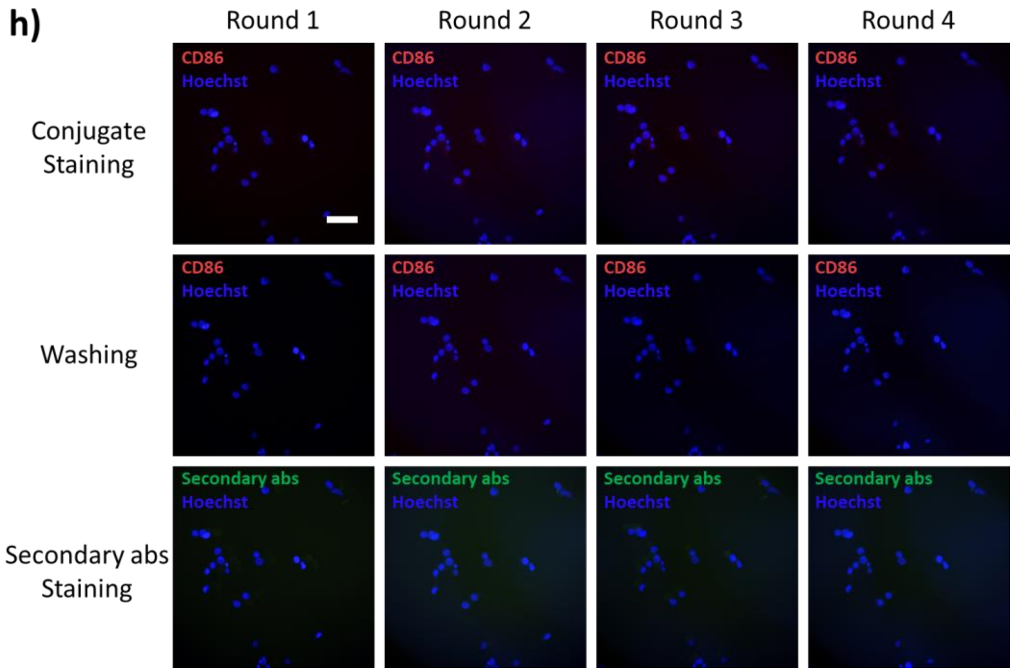


**Figure S6.** Validation of the stability and reproducibility of conjugate staining. The differential N2a cells were treated by 4 consecutive labeling rounds with a) tubulin, b) Lamin B1, c) CREB, d) VGLUT2, e) MAP2, f) phospho-GSK3, g) CD24 and h) CD86 conjugates that were tagged with cDNA-Cy5. For each labeling round, the cells were stained by Cy5 labelled conjugates, washed by a regeneration buffer, and then stained by Alexa Fluor 488 labelled secondary antibodies. Fluorescence images of the cells after each staining and washing step of the 4 consecutive labeling rounds were recorded, respectively. Scale bar = 50 μm.

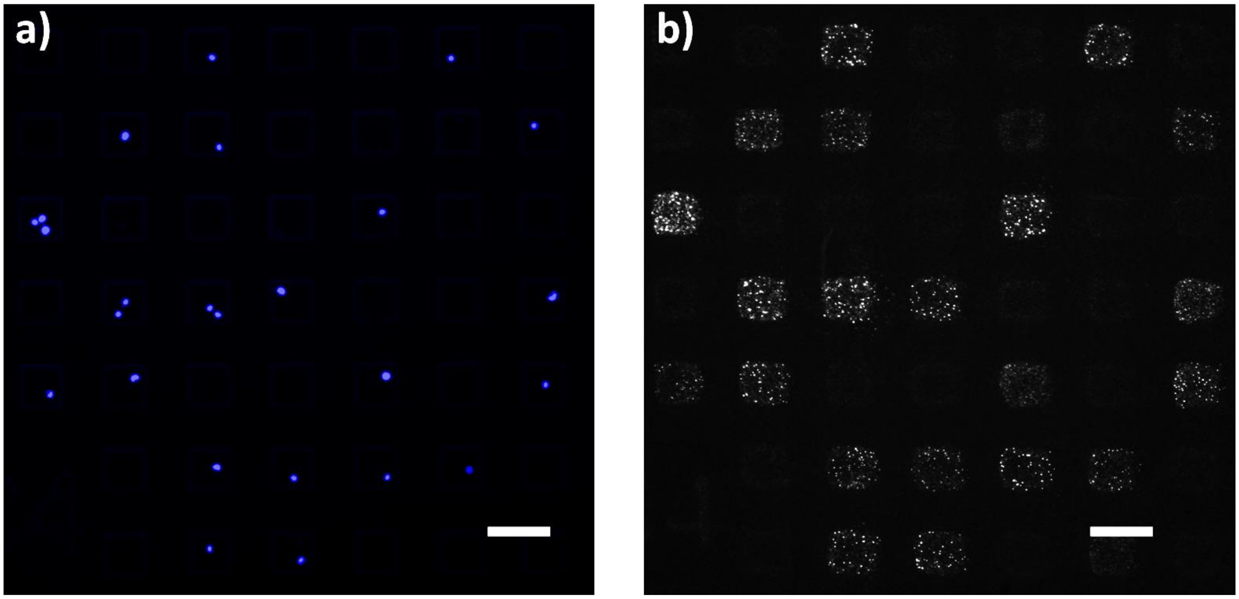


**Figure S7.** Fluorescence images of a) a PDMS microwell chip containing differentiated N2a cells and b) the corresponding protein signal on a MIST array. Cells were labelled with Hoechst dye, and the protein signal was visualized by streptavidin tagged with Alexa Fluor 647 dye. Scale bar = 100 μm.

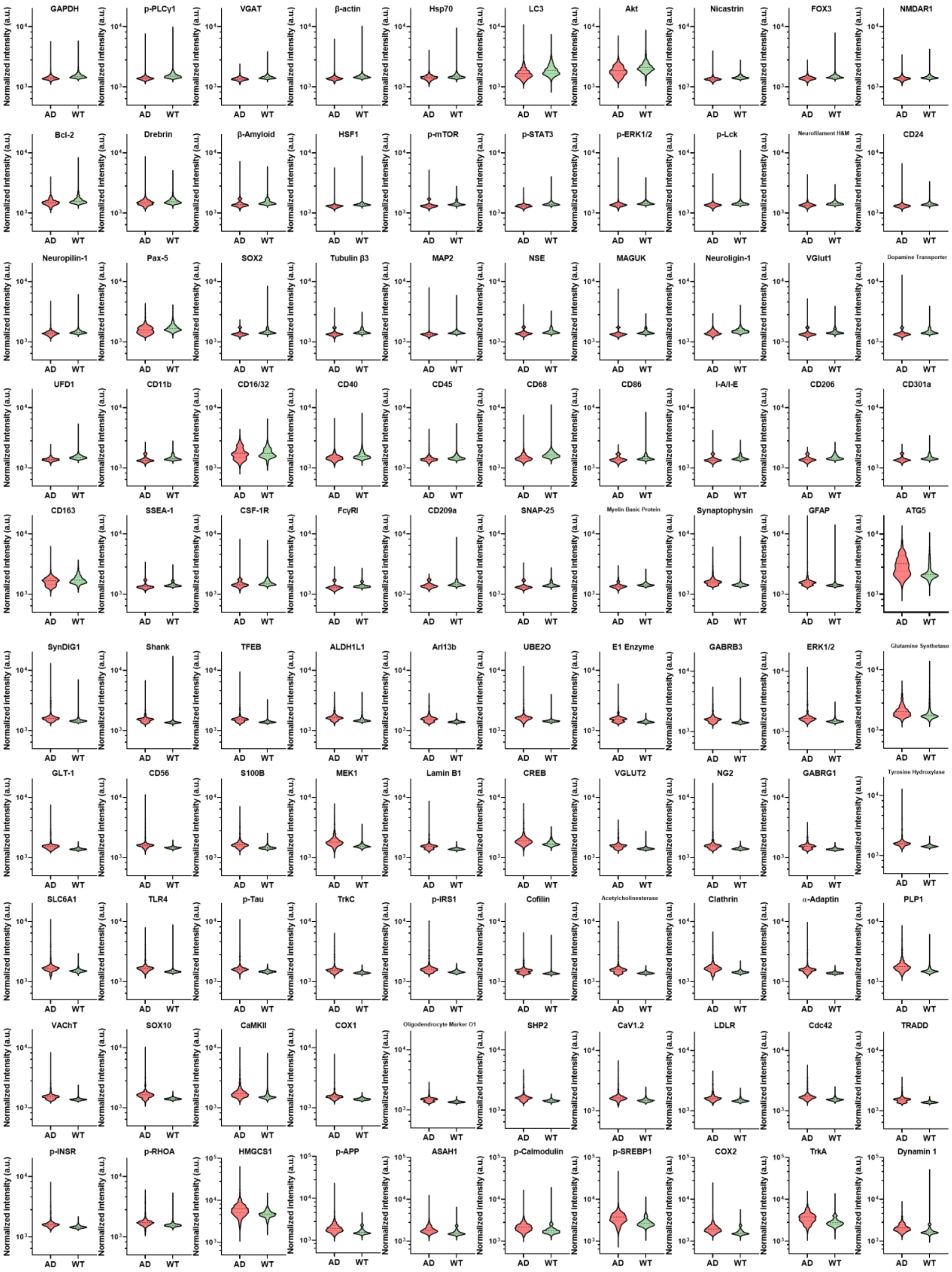

GAPDH
p-PLCγ1
VGAT
β-actin
Hsp70
LC3
Akt
Nicastrin
FOX3
NMDAR1
Bcl-2
Drebrin
β-Amyloid
HSF1
p-mTOR
p-STAT3
p-ERK1/2
p-Lck
Neurofilament H&M
CD24
Neuropilin-1
Pax-5
SOX2
Tubulin β3
MAP2
NSE
MAGUK
Neuroligin-1
VGlut1
Dopamine Transporter
UFD1
CD11b
CD16/32
CD40
CD45
CD68
CD86
I-A/I-E
CD206
CD301a
CD163
SSEA-1
CSF-1R
FcγRI
CD209a
SNAP-25
Myelin Basic Protein
Synaptophysin
GFAP
ATG5
SynDIG1
Shank
TFEB
ALDH1L1
Arl13b
UBE2O
E1 Enzyme
GABRB3
ERK1/2
Glutamine Synthetase
GLT-1
CD56
S100B
MEK1
Lamin B1
CREB
VGLUT2
NG2
GABRG1
Tyrosine Hydroxylase
SLC6A1
TLR4
p-Tau
TrkC
p-IRS1
Cofilin
Acetylcholinesterase
Clathrin
α-Adaptin
PLP1
VAChT
SOX10
CaMKII
COX1
Oligodendrocyte Marker O1
SHP2
CaV1.2
LDLR
Cdc42
TRADD
p-INSR
p-RHOA
HMGCS1
p-APP
ASAH1
p-Calmodulin
p-SREBP1
COX2
TrkA
Dynamin 1
Normalized Intensity (a.u.)
AD
WT

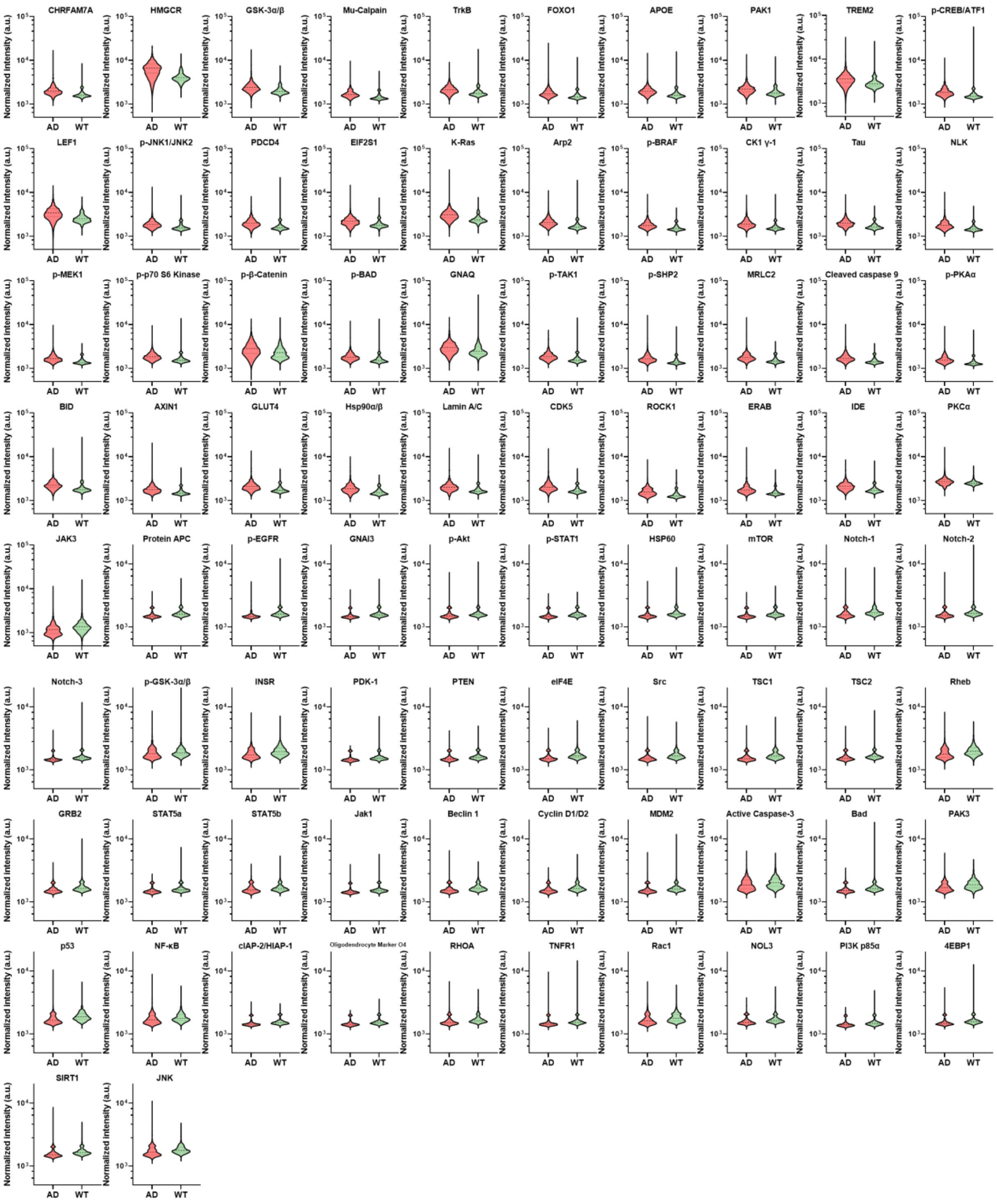


**Figure S8.** Violin plots of the 182 proteins expression level in single cells measured by CycMIST on AD and WT mouse cortex samples.
